# WDR5 regulates epithelial-to-mesenchymal transition in breast cancer cells *via* TGFβ

**DOI:** 10.1101/348532

**Authors:** Punzi Simona, Balestrieri Chiara, D’Alesio Carolina, Bossi Daniela, Dellino Gaetano Ivan, Gatti Elena, Pruneri Giancarlo, Criscitiello Carmen, Carugo Alessandro, Curigliano Giuseppe, Natoli Gioacchino, Pelicci Pier Giuseppe, Lanfrancone Luisa

## Abstract

Even if the mortality rate in breast cancer (BC) has recently decreased, development of metastases and drug resistance are still challenges to successful systemic treatment. The epithelial-to-mesenchymal transition (EMT), as well as epigenetic dynamic modifications, plays a pivotal role in invasion, metastasis, and drug resistance. Here, we report that WDR5, the core subunit of histone H3 K4 methyltransferase complexes, is crucial in coordinating EMT and regulating epigenetic changes that drive metastasis. We show that silencing of WDR5 in BC up-regulates an epithelial signature in triple negative and luminal B like patients by transcriptional repression of mesenchymal genes and reduction of the metastatic properties of these cells. Moreover, we demonstrate that this regulation is mediated by inhibition of the TGFβ signaling both at the transcriptional and post-translational level, suggesting an active role of WDR5 in guiding tumor plasticity upon oncogenic insults, regardless of the pathological BC subtypes.

We therefore suggest that WDR5 inhibition could be a successful pharmacologic approach to inhibit EMT and sensitize breast cancer cells to chemotherapy.

## Introduction

Despite recent advances in BC treatment, women frequently develop resistance to many therapies and die of metastasis (Ginsburg, Bray et al., 2017). Epithelial-to-mesenchymal transition (EMT) is a well-defined developmental program frequently adopted by tumor cells during the metastatic process to gain migratory ability and reach distant organs, losing epithelial cell adhesion and cell-cell contacts, while undergoing cell shape remodeling and cytoskeleton rearrangement (Christiansen & Rajasekaran, 2006, Nieto, 2013). Concurrently, the expression of epithelial markers is inhibited, in favor of an increase in the expression of the mesenchymal genes (Thiery & Sleeman, 2006). The epithelial and mesenchymal states represent two opposite cellular phenotypes that cells reach going through several intermediary phases (Grande, Sanchez-Laorden et al., 2015, Jordan, Johnson et al., 2011, Yu, Bardia et al., 2013).

During EMT, changes in gene expression are crucial for the process to occur. Availability of appropriate transcription factors and degree of openness or closeness of the chromatin play a central role in determining changes in gene expression (Bedi, Mishra et al., 2014). Indeed, the EMT transcriptional program is tightly controlled by both DNA methylation (Cedar & Bergman, 2009) and post-translational histone regulation (Campos & Reinberg, 2009). Even if the epigenetic contribution to the process is not fully understood, several histone modifications are implicated in either inducing or repressing specific sets of EMT genes (Lee & Kong, 2016). The core subunit of the COMPASS-like complex WD repeat-containing protein 5 (WDR5) recognizes the histone H3 amino-terminal tail, thus enabling its methylation on Lys4 (H3K4) (Ford & Dingwall, 2015). WDR5 has a pivotal role in tumor growth and proliferation (Carugo, Genovese et al., 2016, Chen, Xie et al., 2015, Mungamuri, Murk et al., 2013), differentiation (Ang, Tsai et al., 2011, Wang, Han et al., 2017) and metastasis (Lin, Wang et al., 2017, Malek, Gajula et al., 2017, Tan, Chen et al., 2017), showing deregulated expression in different cancers (Carugo et al., 2016, Chen et al., 2015, Cui, Li et al., 2018, Dai, Guo et al., 2015, Ge, Song et al., 2016, Wu, Diao et al., 2018b). Moreover, mesenchymal genes are marked with H3K4me3 by WDR5 in breast cancer upon hypoxia treatment (Wu, Tsai et al., 2011), and in colorectal (Tan et al., 2017), prostate (Malek et al., 2017), and lung ^26^ tumor cells. Despite its proven role in EMT, the causal role of WDR5 in cellular plasticity during EMT still remains elusive.

Transforming growth factor-β (TGFβ) is a multifunctional cytokine involved in a plethora of events regulating EMT in the context of metastasis. Both canonical and non-canonical TGFβ signaling are involved in the regulation of distinct processes as cytoskeleton organization, survival, cell migration and invasion (Micalizzi & Ford, 2009). It has been reported that TGFβ cooperates with other signaling pathways, as well as epigenetic regulators, in the induction of transcription factors, such as the SNAIL, ZEB and TWIST families, driving EMT (Cardenas, Zhao et al., 2016, Chen, Lorton et al., 2017).

Here, we demonstrate that WDR5 is directly involved in the epigenetic regulation of cellular plasticity in BC during EMT and that its inhibition reverts the mesenchymal phenotype mediated by the TGFβ1 response.

## Results

### WDR5 promotes growth of human breast cancer and is a potential therapeutic target

In our previous study, we performed a loss of function shRNA screening in the MCF10DCIS.com (hereafter MCF10DCIS) BC cell line to identify epigenetic targets driving tumorigenesis (D’Alesio, Punzi et al., 2016). Since WDR5 was strongly depleted both in the *in vivo* and *in vitro* screens and it ranked as one of the best candidate in sustaining BC growth (D’Alesio et al., 2016), we first validated its oncogenic role in MCF10DCIS cells. Cells were transduced with two pooled shRNAs to silence WDR5 and a corresponding control (shLuc) (Appendix Fig. S1A) and were either transplanted in NOD/SCID mice to assess *in vivo* tumor growth, or cultured to evaluate the *in vitro* proliferative ability. WDR5 silencing strongly reduced tumor growth *in vivo* (Fig. 1A), as well as proliferation *in vitro* (Fig. 1B), confirming that WDR5 is crucial in sustaining BC growth. Concordantly, we tested the efficacy of the WDR5 small molecule inhibitor, OICR-9429, already proven to be effective in leukemia, pancreatic and lung cancer cells (Carugo et al., 2016, Chen et al., 2017, Grebien, Vedadi et al., 2015). *In vitro* treatment of MFC10DCIS cells with OICR-9429 reduced clonogenicity (Fig. 1C) and proliferation (Fig. 1D) in a dose-dependent manner, similar to that observed by using shRNA.

**Figure 1.**
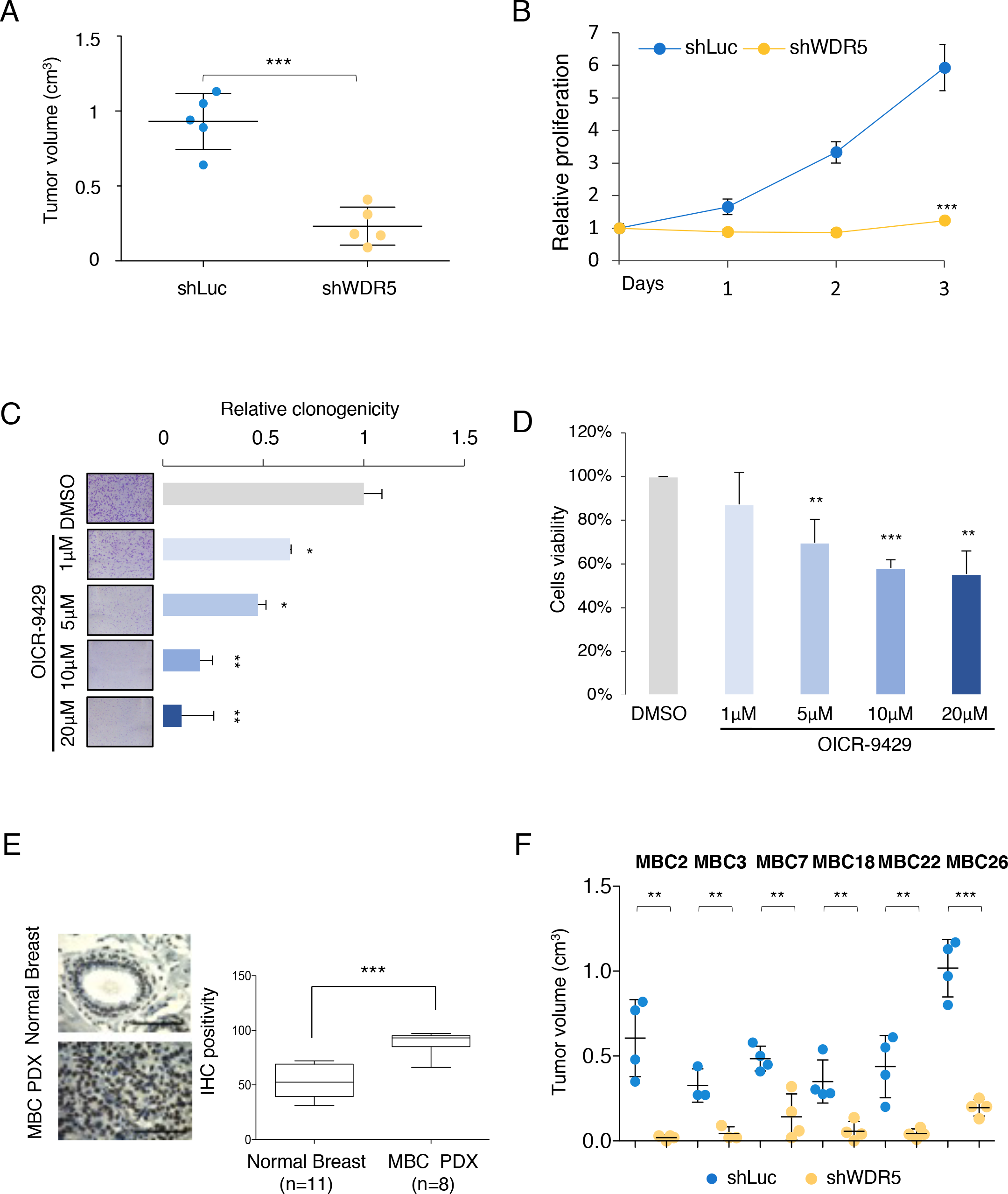
WDR5 is overexpressed in breast cancer and controls tumorigenesis. (A-B) MCF10DCIS cell line was infected to silence WDR5 (shWDR5) or a neutral control (shLuc) and inoculated *in vivo* in immunocompromised mice (n=5 per group) or plated for *in vitro* proliferation assays (n=3). A) Dot plots represent tumor volume in shWDR5 or shLuc MCF10DCIS (mean ± SD - cm^3^). Statistical significance was determined using an unpaired Student *t* test (***: P<0.001). B) *In vitro* relative proliferation values of shWDR5 in MCF10DCIS cells are reported with respect to control shLuc. Statistical significance was determined using an unpaired Student *t* test (***: P<0.001). (C-D) A small molecule inhibitor of *WDR5*, OICR-9429, was used at increasing concentration in MCF10DCIS cells. C) Images and corresponding histograms represented cell clonogenicity expressed as mean ± SD of a biological triplicate after 6 days of treatment. Cells were stained with crystal violet and quantified by using ImageJ analysis. D) Effect of OICR-9429 on MCF10DCIS viability was evaluated after 3 days of treatment and expressed as mean ± SD of a biological triplicate. Statistical significance was determined using an unpaired Student *t* test (*: P<0.05; **:P<0.01; ***: P<0.001). E) WDR5 expression was detected in normal breast tissues (n=11) and Metastatic Breast Cancer (MBC) PDXs (n=8) by IHC. Representative samples of normal and MBC PDX were shown. Scale bar 100μm. Box plots represent WDR5 expression quantified by using Fiji tools for DAB positivity. Statistical significance was determined using an unpaired Student *t* test (***: P<0.001). F) Six MBC PDXs were infected to target control (shLuc) and shWDR5. Transduced cells were transplanted in NSG mice. Dot plots represent tumor volume (mean ± SD – cm^3^) of three to four tumors *per* group. Statistical significances were calculated by applying an unpaired Student *t* test (**: P<0.01; ***: P<0.001).

In order to generate a reliable model to elucidate BC development and treatment opportunities, we created a bank of patient-derived xenografts (PDXs) obtained from liver and lung metastases of BC (MBC) patients who developed resistance to different lines of therapy (Appendix Table S1A). We demonstrated that PDXs recapitulate the phenotypic and molecular features of the tumors (Appendix Table S1B) and we used them as patient surrogates of the metastatic disease to investigate WDR5 expression. Since large evidence show the over-expression of WDR5 in diverse tumor types (Carugo et al., 2016, Chen et al., 2015, Cui et al., 2018, Ge et al., 2016, Wu et al., 2018b), we confirmed that WDR5 expression was significantly higher in MBC PDXs than in the normal mammary gland (Fig. 1E).

Six PDXs, representing triple negative (TN) and luminal B like (LB) BC subtypes, were used for the *in vivo* studies. Two TN (MBC2 and MBC7) and 4 LB (MBC3, MBC22, MBC18 and MBC26) PDXs were independently transduced with two shRNAs WDR5*-*specific in pool or a corresponding control (shLuc) (Appendix Fig. S1B). In parallel, we tested WDR5-silencing effects in highly aggressive TN (SUM149PTand MDAMB468) and LB (HCC1428 and ZR751) BC cell lines (Prat, Karginova et al., 2013) (Appendix Fig. S1C). The reduction of WDR5 expression significantly inhibited tumor growth in MBC PDXs (Fig. 1F) and in all tested cell lines (Appendix Fig. S1D). *In vitro* proliferation was also significantly impaired by shWDR5 (Appendix Fig. S1E), suggesting that WDR5 may be involved in tumorigenesis, independently of the BC subtypes.

### WDR5 expression promotes breast cancer metastasization

Despite the fact that WDR5 has been reported to promote migration *in vitro* and metastasization *in vivo* in colon and prostate cancer cell lines (Malek et al., 2017, Tan et al., 2017), its role in BC is still unproven. We therefore compared the metastatic ability of shLuc and shWDR5 MDA-MB-231 cells by performing functional assays both *in vivo* and *in vitro*. Cells were first transduced with the Luciferase gene and then with either the WDR5 shRNA or the neutral control (SCR) and transplanted in the 4^th^ mammary gland of NSG mice. Tumors were resected at the same volume in the control and WDR5-silenced groups (Appendix Fig. S2A) and mice were weekly tested for metastasis formation by bioluminescence assay. MDA-MB-231 metastasized to lung and axillary lymph nodes in SCR (Fig. 2A, upper panel) transplanted mice, while WDR5 silencing totally inhibited (3/5), or significantly reduced (2/5), the number and size of the metastatic foci (Fig. 2A, lower panel), as confirmed by the histological staining of the metastatic organs (Appendix Fig. S2B).

**Figure 2.**
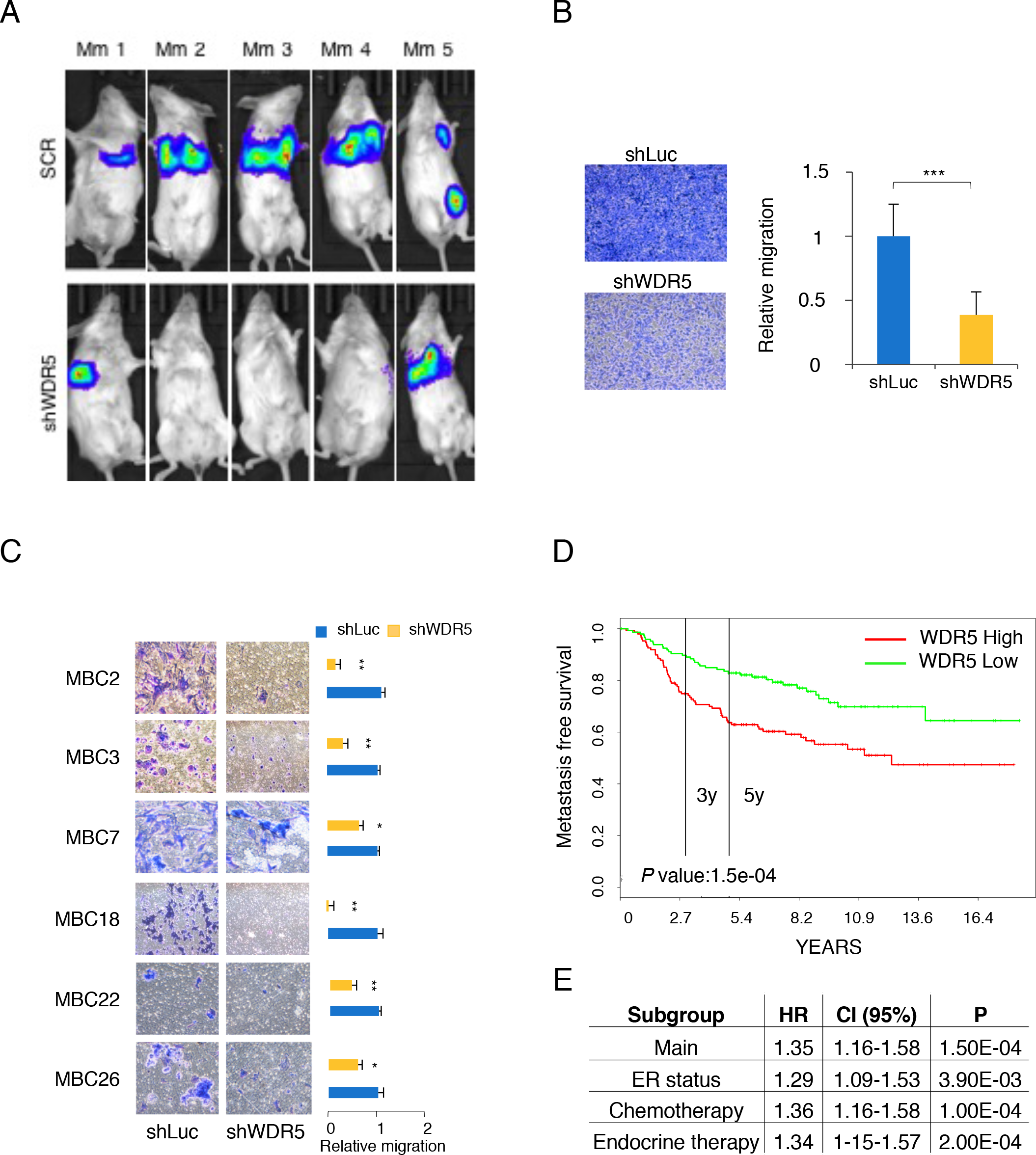
WDR5 silencing significantly affected breast cancer metastasization. Effect of WDR5 silencing was studied in MDA-MB-231 as breast cancer metastatic model. A) MDA-MB-231 cells were infected to express Luciferase, transduced to silence WDR5 or a control vector (SCR) and then transplanted in NSG mice (n=5 per group). Metastasis detection at distant organs was performed by using bioluminescence technique *in vivo* after resection of primary tumors. B) Relative migration was performed on transwells (n=3), stained by crystal violet and represented as ratio of shWDR5 and shLuc values. Statistical significance was determined using an unpaired Student *t* test (***: P<0.001). C) Cells from MBC PDXs were used for *in vitro* migration study in duplicate. Relative migration (mean ± SD) was calculated by ImageJ analysis on Crystal violet staining positivity and expressed as ratio of silenced versus control values. Statistical significances were calculated by applying an unpaired Student *t* test (*: P<0.05; **: P<0.01). D) Metastasis free survival in breast cancer patients (n=295) was calculated in WDR5-low and WDR5-high groups divided at median of gene expression. Statistic comparison was performed using log rank test. E) WDR5 expression was considered with respect to estrogen status, or chemotherapy or endocrine therapy and Hazard Ratio (HR), confidence interval (CI)(95%) and P value were calculated.

Modified Boyden chamber assay consistently demonstrated that WDR5 silencing led to a substantial reduction of the MDA-MB-231 cell migration (Fig. 2B), as well as in 6 MBC PDXs (Fig. 2C) and in MCF10DCIS (Appendix Fig. S2C) and the TN and LB cell lines (Appendix Fig. S2D) mentioned above. Concordantly, wound healing assay (Appendix Fig. S2E, S2F), performed in shWDR5 MDA-MB-231 cells, displayed a significant impairment of cell migration, as well. Overall these data suggest that WDR5 impacts on the ability of BC cells to migrate and metastasize to distant organs, and that its over-expression may be predictive of metastatic disease. To this aim, we analyzed WDR5 expression and Metastasis Free Survival (MFS) in a NKI data set of 295 patients, showing that higher WDR5 expression was associated with worse MFS (Fig. 2D), and that the prognostic role of WDR5 was independent of the hormonal and therapeutic status (Fig. 2E).

### WDR5 silencing inhibits EMT in breast cancer

The ability of BC cells to metastasize has been correlated with the acquisition of an EMT phenotype (Chaffer, San Juan et al., 2016). Further, previous works suggested that WDR5 might regulate metastasis formation by inducing EMT in various cancers (Chen et al., 2017, Lin et al., 2017, Malek et al., 2017, Tan et al., 2017, Wu et al., 2011, Wu, Xiang et al., 2018a). We tested whether WDR5 silencing could reduce the migratory ability of BC cells by directly inhibiting EMT. As maintenance of the epithelial phenotype requires the coordinated expression of a set of genes directly implicated in cell adhesion and cytoskeletal polarization, we evaluated the transcriptional profile of control (shLuc) and shWDR5-silenced PDXs cells (2 TN and 3 LB) by RNA-seq. For each PDX we performed pairwise analysis to identify differentially expressed genes (DEGs) (Fig. 3A). In order to exclude individual specificities, possibly due to the intra-and inter-tumor heterogeneity of BC (Polyak, 2011), we considered a Golden Gene Set (GGS) of 161 down-and 92 up-regulated genes in common among at least 2 PDXs (Fig. 3B, Appendix Table S2). Gene Ontology (GO) analysis of GGS is shown in Figure 3C. Top ranking categories were related to cell polarization and adhesion, consistently reflecting changes in cell shape and cytoskeleton rearrangement that affect cell motility (Fig. 3C and Appendix Table S2). The inhibition of migratory cell ability may be in part explained by down-regulation of MAPRE1, a master regulator of microtubules (Rovini, Gauthier et al., 2013) and SDCBP, related to polarized distribution of F-actin (Das, Pradhan et al., 2018). Moreover, HSPA1B and PPME1 over-expression has been associated with poor survival of patients with ovarian, gastric and lung cancers (Jakobsson, Moen et al., 2015, Li, Han et al., 2014), suggesting that their down-modulation may reduce tumor progression. Concordantly, actin cytoskeleton rearrangements were detected in shLuc and shWDR5 MCF10DCIS cells. Immunofluorescence analysis revealed that filamentous actin (F-actin) was assembled into actin stress fibers in control cells, while in shWDR5 cells F-actin predominantly organized in cortical bundles tightly associated with cell-cell adhesions (Fig. 3D), reminiscent of an epithelial-like phenotype. We also investigated the adhesion properties of WDR5-silenced MCF10DCIS cells on a panel of extracellular matrices (collagen, laminin, fibronectin, matrigel), showing that cell adhesion is increased in shWDR5 cells both in term of cell number (Fig.3E and Appendix Fig. S3A) and cell spreading (Fig.3E and Appendix Fig. S3B), thus suggesting a tighter cell anchoring to the matrix.

**Figure 3.**
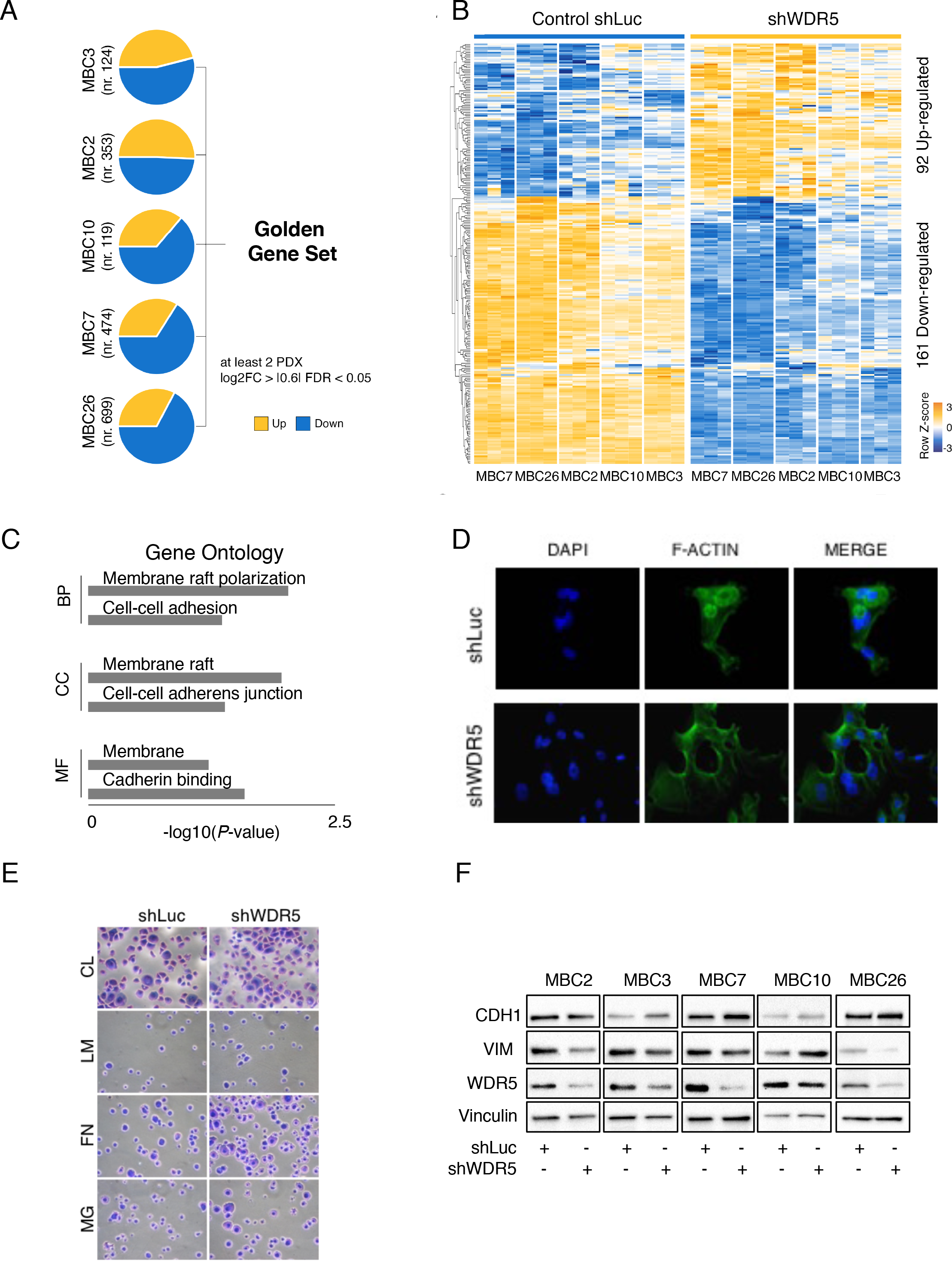
WDR5 silencing inhibits EMT in breast cancer. RNA-seq was performed in 5 PDXs upon WDR5 silencing. A) Pie charts showed differentially expressed genes (DEGs) obtained by pairwise analysis in each PDX comparing shLuc and shWDR5 MBC PDXs cells. Genes were identified as DEGs when the following criteria were met: log_2_ fold-change (FC) ≥ |0.6|, false discovery rate (FDR) < 0.05 and expression > 0.5 RPKM. DEGs in common at least in two PDXs defined a Golden Genes Set (GGS). B) Heatmap showed normalized expression values after removing batch effects using 253 DEGs belonging to GGS. Blue and orange colors indicate down-and up-regulated genes, respectively. C) Representative Gene Ontology terms enriched in DEGs are shown. For complete list refer to Appendix Table S2. D) Morphological changes by actin cytoskeleton remodeling in MCF10DCIS breast cancer cell line due to *WDR5* silencing were evaluated by immunofluorescence technique (n=3). Images show F-actin, DAPI and merged staining in shLuc and shWDR5 cells. E) Adhesion to a panel of substrates (collagen - CL, laminin - LM, fibronectin - FN, matrigel - MG) in shLuc or shWDR5 cells was considered (n=3). Cells were stained by crystal violet and adhesion percentage was calculated by ImageJ and expressed as ratio of relative cells area of shWDR5 versus shLuc. Statistical significances among areas were calculated by applying an unpaired Student *t-*test (***: P<0.001). F) Effect of WDR5 silencing on E-cadherin (CDH1) and Vimentin (VIM) expression was detected by western blot in PDXs used for transcriptomic analysis. Vinculin was used as normalizer.

We further characterized the expression of epithelial and mesenchymal markers in MBC PDXs in response to WDR5 silencing. WDR5 knockdown increased protein levels of E-cadherin (CDH1), a prominent biomarker of the epithelial state, while reduced the mesenchymal marker Vimentin (VIM) (Fig. 3F), suggesting that WDR5 may have a direct role in phenotype reversion. Moreover, the majority of genes acting as mesenchymal inducers or epithelial repressors, i.e. CDH2, TWIST1, SNAI1, SNAI2 and ZEB1, were transcriptionally down-modulated upon WDR5 silencing (Appendix Fig. S3C). Since, the transition from a mesenchymal to an epithelial state with drug treatment can be used to enhance lethality and eradicate epithelial cells, as well as a strategy to sensitize cells to chemotherapy (Meidhof, Brabletz et al., 2015), we used MCF10DCIS and MDA-MB-231 to test the effect of WDR5 inhibition on lethality and sensitization to paclitaxel (PTX). PTX administration, upon WDR5 silencing, significantly reduced MCF10DCIS cells viability after 72h of exposure with respect to WDR5-untreated cells (Appendix Fig. S3D). Interestingly, MDA-MB-231, that showed higher resistance to PTX, displayed significantly enhanced cell death also at low PTX doses upon WDR5 silencing (Appendix Fig. S3E).

Conversely, over-expression of WDR5 caused changes in cell morphology, leading to an elongated spindle shape of the cells (Appendix Fig. S4A), accompanied by the induction of mesenchymal gene expression (Appendix Fig. S4B), and increased tumorigenicity of MCF10DCIS cells *in vivo* (Appendix Fig. S4C). Overall, these observations suggest that WDR5 regulates the EMT program at transcriptional level in breast cancer cells and that its silencing can inhibit the mesenchymal phenotype.

### WDR5 mediates epigenetic regulation of EMT

The EMT program is governed by a series of complex epigenetic interactions of transcription factors with protein complexes, whose mechanisms of action in driving plasticity during transition is still debated (Skrypek, Goossens et al., 2017). WDR5, as the core component of histone H3K4 methyltransferase complexes, is responsible of the methylation pattern of many EMT genes (Chen et al., 2017, Wu et al., 2011). We first analyzed WDR5 occupancy at the promoter regions of genes in MCF10DCIS cells by chromatin immunoprecipitation (ChIP) and sequencing. We found that around 80% (4413/5794) of the peaks were located proximal to the Transcription Start Sites (TSS) (± 3000 bp from TSS), confirming that WDR5 regulates its targets mainly at the promoter level (Fig. EV 1A). KMT2A/MLL1 and ERBB2, already known interactors of WDR5 (Ford & Dingwall, 2015, Mungamuri et al., 2013), are among these genes, thus validating our analysis. We also showed that TGFβ1, SMAD2 and SMAD3 and some members of the

PI3K complex displayed a peak of WDR5 on their promoters, suggesting that WDR5 may be involved in multiple signaling regulation (Table EV 1). We performed additional RNA-seq analysis in shLuc and shWDR5 MCF10DCIS cells, and we detected 1’315 down-and 919 up-regulated DEGs. Diverse epithelial identity genes, keratins, genes regulating tight junctions and focal adhesions, as well as cell-cell adhesion molecules were found up-regulated (Fig. EV 1B). On the contrary, genes commonly involved in promoting mesenchymal phenotype and metastasis (i.e. TGFβ1, FOXQ1, ERBB3, S100A14, KIF4, CCR1, HAS2, PAK1 and TIAM1) were down-regulated (Fig. EV 1B), thus suggesting that WDR5 knockdown may sustain an epithelial phenotype. We therefore assessed H3K4me3 intensity at the promoter level of the expressed genes (± 1’500 bp from TSS) (Table EV 3) in MCF10DCIS cells, and we observed a global reduction of intensity in shWDR5 with respect to shLuc (*P*= 4.5E-9) (Fig. 4A). Remarkably, a statistical significant reduction in H3K4me3 intensity was observed at the promoter of the down-regulated (*P* = 1.2E-15), but not of up-regulated genes (Fig. 4A), confirming that the main effects due to WDR5 knockdown were observed in the down-modulated genes and that H3K4me3 signals, which associate with active and poised TSS, correlated with WDR5-dependent gene expression. We also determine level of H3K4me3 intensity at WDR5 peaks in the DEGs in MCF10DCIS cells. Among 907 hypo-methylated regions associated to gene promoters, 260 were found to show both differential expression and H3K4 trimethylation level (238 showed the same direction of alteration). Importantly, we detected 62 genes where WDR5 peaks are close to the TSS of the regulated genes (Fig. EV 1C). Among the top interesting ones we listed TGFβ1 (Fig. 4B), suggesting that WDR5 is essential for H3K4 methylation on TGFβ1 promoter and may be prominent for its activation. On the other side, among 698 hyper-methylated regions, 58 were also differentially expressed and 51 showed concordance in H3K4me3 increase. Only 5 out of 58 regions were directly bound by WDR5 (Fig. EV 1C). Interestingly, TGFβ signaling scored as one of the most significantly enriched biological functions in MBC PDXs (Appendix Table S2). The modulation of TGFβ-dependent pathways includes down-regulation of many oncogenes (Fig. 4C): (i) TGFβR2, as TGFβ type II receptor sustains metastasis of high-grade BC tumors (Tang, Vu et al., 2003); (ii) FURIN, a proprotein convertase that promotes the malignant phenotype of cancer cells (Jaaks & Bernasconi, 2017) and stimulate BC invasion (Li, Chu et al., 2018); (iii) GCNT2, that modulates EMT by modification of E-cadherin glycosylation (Zhang, Meng et al., 2011);
(iv) PARD6, crucial for cell polarization (Mu, Zang et al., 2015) and (v) LTBP2, the Latent TGFβ-Binding protein, whose knock down inhibits tumorigenesis in thyroid carcinomas (Wan, Peng et al., 2017). Conversely, the tumor suppressor TGFβR3 was up-regulated in WDR5 silenced cells (Fig. 4C and Appendix Table S2), consistent with its role in suppressing metastasis formation in clear-cell renal cell carcinoma (Nishida, Miyazono et al., 2018). Finally, Ingenuity Pathway Analysis (IPA) confirmed that TGFβ1 resulted the top-ranking among genes or complexes that regulate EMT in breast cancer (HDAC3, ERBB2, HIF1A, SMAD7)(Ingthorsson, Andersen et al., 2016, Papageorgis, Lambert et al., 2010, Wu et al., 2011), as well as to be involved in tumor progression and metastatic dissemination (i.e. PI3K, E2F4, VEGFA and STAT3) (Beshiri, Holmes et al., 2012, Das et al., 2018, Jang, Kim et al., 2017, Tan et al., 2017). When transcriptomic profiles of the GGS of the PDXs and of the MCF10DCIS cells have been intersected, 99 down-regulated and 40 up-regulated genes were found in common (Fig. 4E). In details, we found that 6 out of 17 genes included in the TGFβ1-pathway have a role in sustaining cell migration in a number of cancer cells and were concordantly down-modulated in the majority of the MBC PDXs (Fig. 4F), as well as in MCF10DCIS (Fig. EV 1D). Interestingly, CEMIP (Cell Migration Inducing Hyaluronan Binding protein) expression regulates invasion and metastasis in colon cancer (Evensen, Li et al., 2015), HSPA1B (Heat Shock 70kDa Protein 1B) in ovarian cancer (Jakobsson et al., 2015), AIF1L (Allograft Inflammatory Factor 1 Like) determines cytoskeleton rearrangement (Lu, Ye et al., 2017), and PNP (purine nucleoside phosphorylase) affects prostate cell migration (Kojima, Chiyomaru et al., 2012), while the up-regulation of ECE1 (Endothelin Converting Enzyme 1) (Smollich, Gotte et al., 2007), GBP1 (Guanylate Binding Protein 1) (Pedersen, Hood et al., 2017), HMGCR (3-hydroxy-3-methylglutaryl-CoA reductase) (Singh, Yadav et al., 2015), PARD6A (par-6 family cell polarity regulator alpha) (Viloria-Petit, David et al., 2009) is linked to poor outcome and increased metastatic potential in BC.

**Figure 4.**
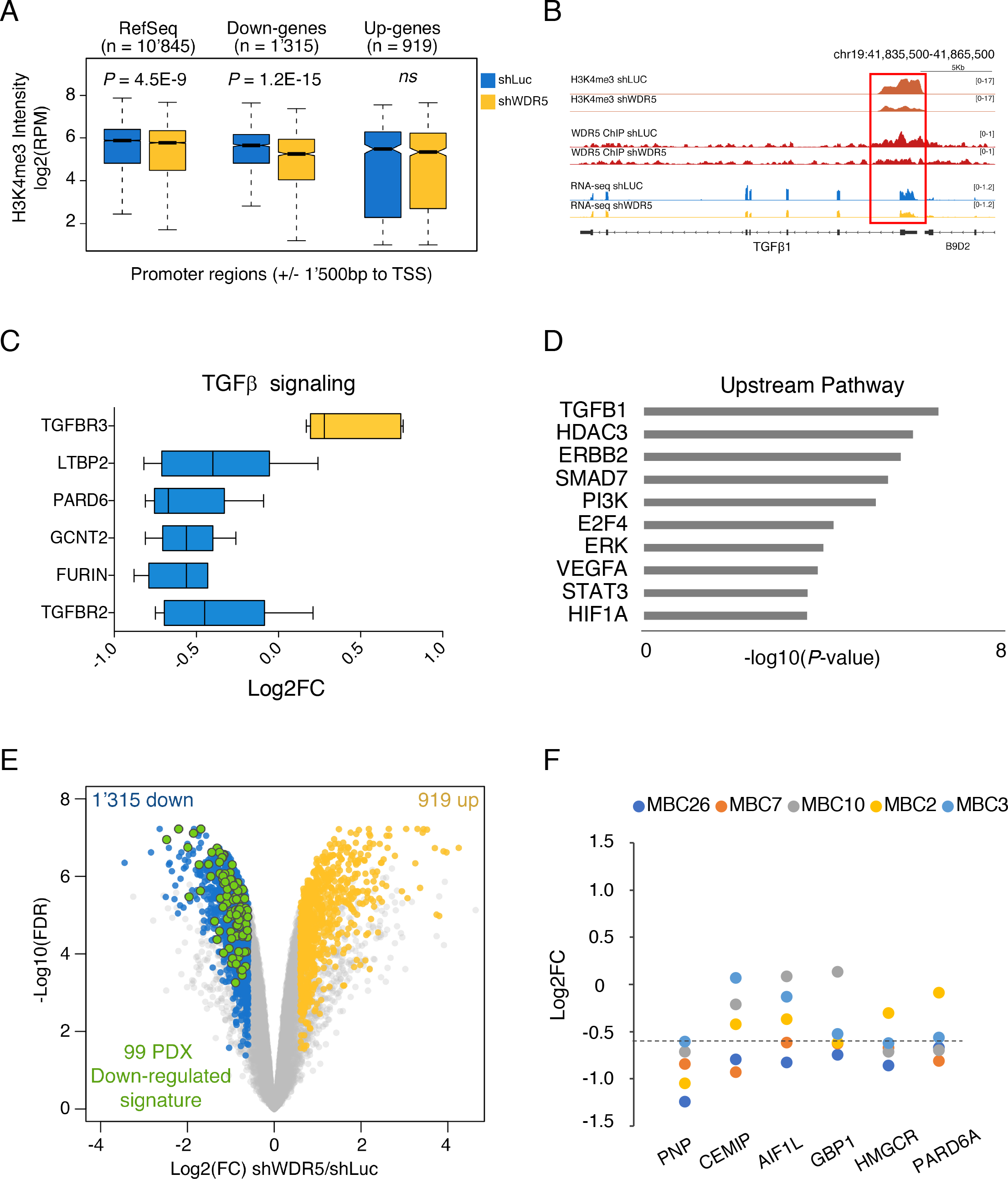
WDR5 regulates its targets by histone methylation. A) Box plots show H3K4me3 signal intensity in shLuc and shWDR5 MCF10DCIS cancer cell lines at gene promoters. Central values represent the median, the bars the 25^th^ and 75^th^ percentile and the dashed lines the lower and upper whiskers. P-value was obtained from a Wilcoxon rank-sum test. B) Snapshot represents WDR5 and H3K4me3 binding peaks identified by ChIP-seq analysis and RNA signal of TGFβ1 gene in a representative replicate of shLuc and shWDR5 MCF10DCIS cells. C) Box plots show Log2FC values for each gene belonging to TGFβ pathway highlighted by Gene Ontology analysis in 5 MBC PDXs. D) Representative IPA upstream pathways enriched in Golden Genes Set of PDXs are shown. For complete list refer to Appendix Table S2. E) Volcano plot shows Differentially Expressed Genes (DEGs) in shWDR5 MCF10DCIS cancer cell line. The log2 expression fold change (FC) is shown on the horizontal axis and −log10 of the FDR is shown on the vertical axis. Up-regulated genes (FDR < 0.05 and log2 FC > 0.6) are highlighted in orange while down-regulated genes in blue (FDR < 0.05 and log2 FC < −0.6). The genes indicated in green are also down-regulated in the PDX set. F) Dot plots represent Log2FC values for each gene in MBC PDXs. Dashed line represents threshold imposed on Log2FC to determine DEGs.

### WDR5 mediates the EMT response to TGFβ1

TGFβ1 is a pleiotropic cytokine that exerts several different functions in the cell, whose dysregulation mainly impacts on cancer progression and regulation of immune surveillance (David & Massague, 2018). Since our finding supported that WDR5 is required for EMT, as well as for TGFβ1 signaling during BC progression, we treated shLuc and shWDR5 MCF10DCIS cells with recombinant human TGFβ1. Then, we tested the relative expression of the mesenchymal genes known to be induced by TGFβ1, namely CDH2, TWIST1, SNAI1, SNAI2, ZEB1 (Micalizzi & Ford, 2009), also showed to be regulated in our PDXs. TGFβ1 dramatically increased the expression of these genes without altering WDR5 levels (Appendix Fig. S5A), while the WDR5 silencing strongly reduced TGFβ1-dependent transcription of these targets (Figure 5A). As shown in Figure 5B and Appendix Fig. S5B, cell migration was stimulated by TGFβ1 and restored to control levels in presence of WDR5 silencing, thus confirming that TGFβ1-induced migration and EMT is WDR5 dependent. Since it has been reported that TGFβ1 promotes EMT by a combination of canonical

**Figure 5.**
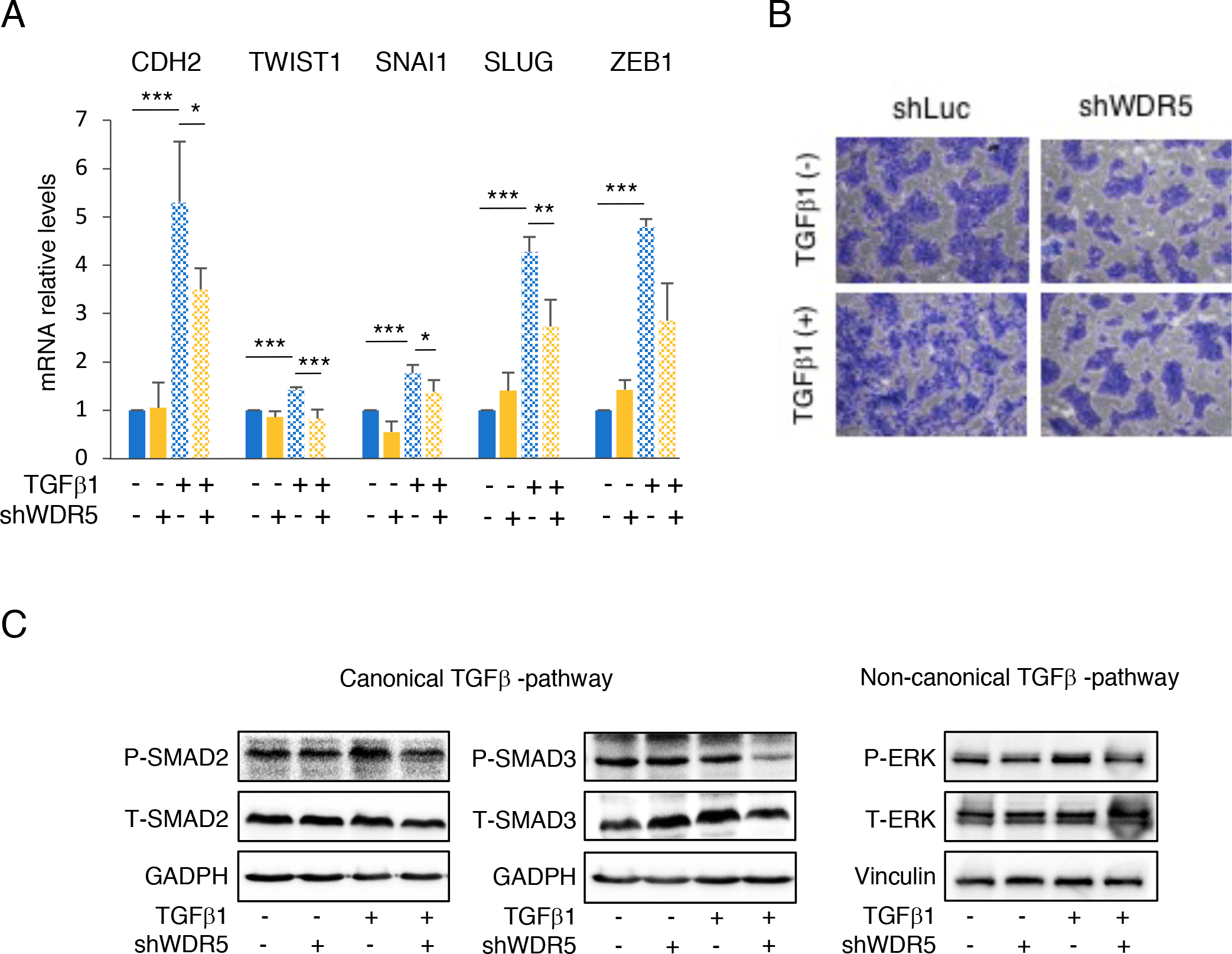
EMT and metastasis are regulated by TGFβ1-WDR5 axis in breast cancer. A) Relative expression of CDH2 and TWIST1, SNAI1, SNAI2 and ZEB1 transcription factors in MCF10DCIS breast cancer cell line was analyzed by RT-qPCR. Cells were treated with (+) or without (−) 5 ng/mL of TGFβ1 for 2 days and with (+) or without (−) shWDR5. Statistical significance was determined using an unpaired Student *t* test (*: P<0.05; **: P<0.01; ***: P<0.001) B) Migration of MCF10DCIS cells treated with (+) or without (−) TGFβ1 and with (+) or without (−) shWDR5 is shown. C) Immunoblots for total-(T) or phosphorylated-(P) SMAD2 and SMAD3, and (D) T-ERK and P-ERK were performed in MCF10DCIS cells treated with (+) or without (−) TGFβ1 and with (+) or without (−) shWDR5. GADPH or Vinculin were used as normalizers.

SMAD-dependent and non-canonical SMAD-independent transcriptional events (through the ERK1/2 pathway) (Felipe Lima, Nofech-Mozes et al., 2016), we wanted to confirm whether the TGFβ1 downstream mediators SMAD2/3 and ERK1/2 were also prominent for WDR5 regulation of TGFβ1 signaling. In our system, SMAD2, but not SMAD3, was phosphorylated by TGFβ1, while silencing of WDR5 reduced the phosphorylation of both substrates (Fig. 5C). Also, phosphorylation of ERK1/2, induced by TGFβ1, was inhibited by shWDR5 (Fig. 5D), thus confirming that various integrated signaling pathways are required for the regulation of WDR5 on TGFβ1.

### Discussion

A extensive body of literature supports the essential role of WDR5 in targeting histones to epigenetically regulate cancer progression (Carugo et al., 2016, Chen et al., 2017, Ge et al., 2016, Mungamuri et al., 2013). TGFβ signaling and EMT are strictly interconnected pathways mediated by many key factors including epigenetic and transcriptional regulation (Dumont, Wilson et al., 2008). In this study we report that WDR5 is an essential gene in breast cancer development and progression that works as a crucial intermediate in coupling TGFβ pathway and EMT.

It has been previously shown that WDR5 expression is frequently deregulated in leukemia, colon, pancreatic and prostate cancer (Carugo et al., 2016, Chen et al., 2015, Cui et al., 2018, Ge et al., 2016, Tan et al., 2017). Here, we directly link WDR5 expression to breast cancer progression and metastasis by using a panel of cell lines and PDX models of metastatic breast cancer resistant to different lines of therapy. For these patients offering new therapeutic strategies is mandatory. For *in vitro* and *in vivo* studies, we made use of the highly plastic MCF10DCIS breast cancer cell line, known to be able to recapitulate all the stages of breast cancer progression, from *in situ* growth to metastasis formation (Barnabas & Cohen, 2013, D’Alesio et al., 2016). MCF10DCIS cells appear to be the most attractive model to study the molecular mechanisms responsible of the epigenetic regulation in the transition from the epithelial to the mesenchymal state and *viceversa*.

WDR5 knockdown was able to reduce tumor growth in LB and TN breast cancers, concordantly with the role of WDR5 in regulating tumorigenesis (Carugo et al., 2016, Chen et al., 2015, Mungamuri et al., 2013). We offered direct evidences of the involvement of WDR5 in breast cancer progression by performing a meta-analysis on metastasis free survival in breast cancer patients, as well as by using *in vivo* MDA-MB-231 cells and *in vitro* migratory PDXs.

By employing gene expression profiling of differentially expressed genes in PDXs and breast cancer cell line models, we found that WDR5 transcriptionally regulates an epithelial signature and induce cell adaptation to signaling events. Among these, TGFβ regulation is affected by down-modulation of TGFβ receptors and downstream targets genes that were reported to control cell migration in tumors (Evensen et al., 2015, Jaaks & Bernasconi, 2017, Kojima et al., 2012, Lu et al., 2017, Mu et al., 2015, Pedersen et al., 2017, Singh et al., 2015, Smollich et al., 2007, Tang et al., 2003, Viloria-Petit et al., 2009, Wan et al., 2017, Zhang et al., 2011) (Fig. 6A). By examining the genome wide WDR5 occupancy, we show that WDR5 directly binds to a subset of gene promoters, among which we highlighted TGFβ1, suggesting that WDR5 activates TGFβ oncogenic pathways known to be involved in tumorigenesis and metastasis (Felipe Lima et al., 2016). Indeed, transcriptional regulation of WDR5 targets occurs through recruitment of KMT2A in the COMPASS-like complex, and subsequent methylation of H3K4 (Alicea-Velazquez, Shinsky et al., 2016). Our data suggest that a similar mechanism is in place also in breast cancer, as WDR5*-*dependent, down-regulated genes are hypomethylated in WDR5 silenced cells. TGFβ1 signaling has been widely associated with the EMT phenotype (David & Massague, 2018). Mimicking microenvironmental sources, TGFβ1 administration maintains TGFβ signaling overactive and sustains invasive properties of BC cells by altering many EMT transcriptional drivers, including TWIST1, SNAI1, SNAI2 and ZEB1. It is well known that these transcription factors may recruits chromatin modifiers (HDAC1, HDAC2, LSD1, PRC2 complex) by activating dynamic chromatin control loop of their targets (Herranz, Pasini et al., 2008, Lin, Ponn et al., 2010, Peinado, Ballestar et al., 2004) and contributing to the plasticity during transition. EMT induced by exogenous TGFβ was significantly impaired by WDR5 silencing, by reducing transcription of many EMT genes and bringing the cells to an intermediated metastable state. This effect also reflects the flexibility of these cells to undergo EMT, or revert the mesenchymal phenotype, leading to metastasis (Bedi et al., 2014) (Fig. 6B). Finally, WDR5 is instrumental in allowing TGFβ canonical (SMADs) and non-canonical (ERK1/2)-dependent activation since the impact on phosphorylation strongly implies a direct involvement of WDR5 in the post-translational modulation of the TGFβ machinery.

Overall, we suggest a key role of WDR5 in cell program by driving tumor plasticity upon oncogenic insults. Reversion of the EMT program by inhibition of WDR5 represents a captivating therapeutic option, in particular for drug-resistant patients. In fact, we have shown that WDR5 inhibition can sensitize breast cancer cells to paclitaxel even at low doses, suggesting a promising way of counteracting the onset of resistance to second line therapies. These observations are encouraging for further clinical investigation and suggest that WDR5 inhibition could be a novel therapeutic approach to prevent tumor metastasis in breast cancer.

**Figure 6.**
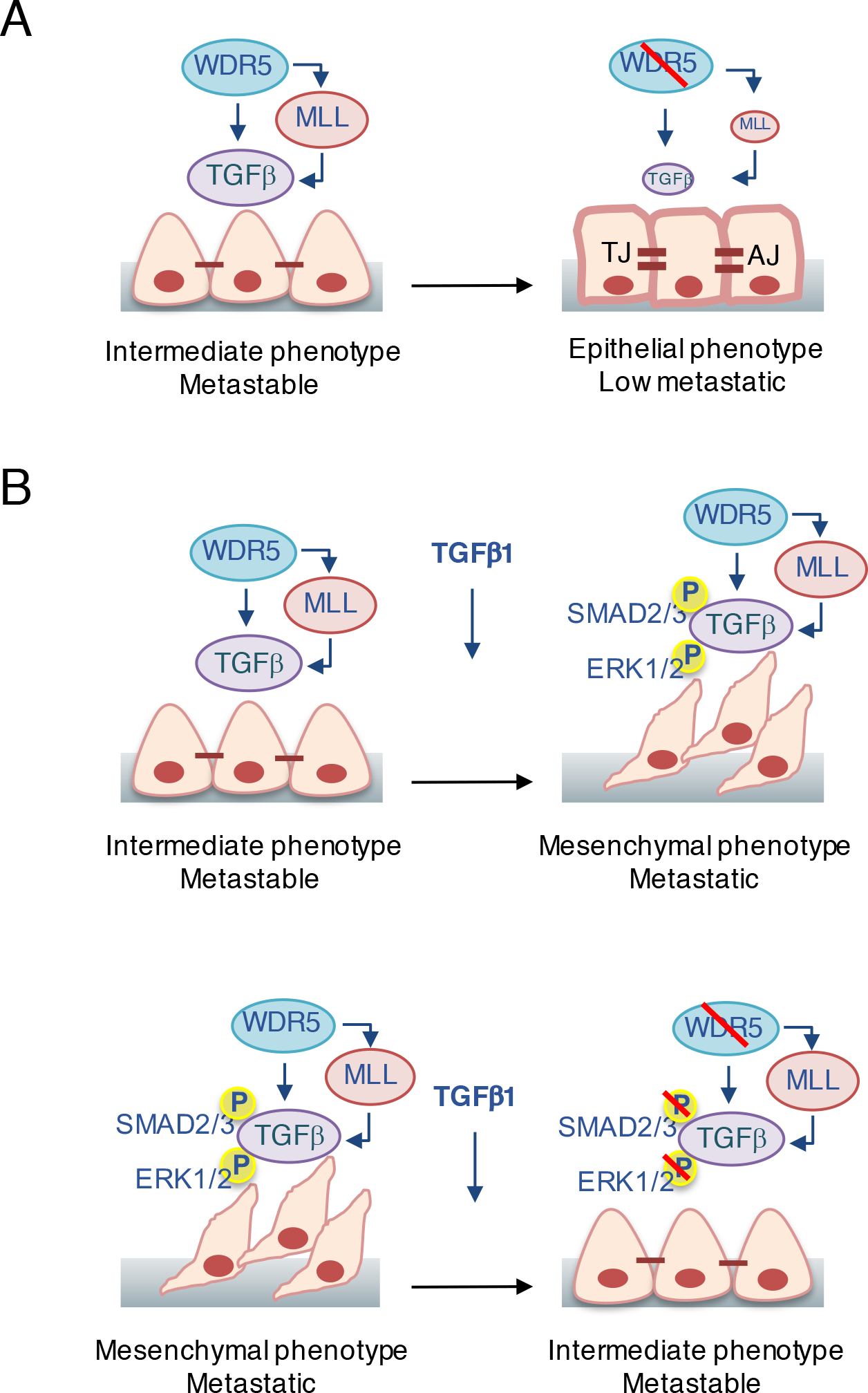
Molecular mechanism of WDR5-TGFβ1 axis in the regulation of EMT in breast cancer cells. A) In breast cancer, silencing of WDR5 is able to induce cell differentiation from an intermediate mesenchymal toward an epithelial–like phenotype through transcriptional down-regulation of the endogenous levels of TGFβ1 and the contextual down-regulation of H3K4me3 intensity at its promoter. The epithelial trait is supported by the up-regulation of keratins, tight junction (TJ), adherent junction (AJ), remodeling of cytoskeleton and down-regulation of mesenchymal genes. Interestingly, WDR5-dependent reprogramming switches cells from metastable to low metastatic. B) Exogenous TGFβ1 perturbation, frequently observed during metastasization, is able to induce EMT and the metastatic phenotype, by phosphorylation of TGFβ-canonical (SMAD2/3) and non-canonical (ERK1/2) complexes (upper panel). On the contrary, WDR5 silencing partially abrogates phosphorylation, inhibits EMT genes transcription and reduces cell migration, thus bringing breast cancer cells to the previous metastable state (lower panel).

## Methods

### PDX tissue bank generation

Patients enrolled in the study were selected on the basis of highly aggressive metastatic disease diagnoses (Luminal B and Triple Negative subtypes) and resistance to different lines of therapy. Biopsies from liver and lung were transplanted in Matrigel (Corning #356231) orthotopically in the 4^th^ mammary gland of female NSG mice. Tumors were monitored weekly and serially passaged in NSG after tissue digestion (see “PDX culture”). Tumors were characterized by IHC on the basis of the prognostic clinical markers, i.e. Estrogen, Progesterone, HER2 and Ki67, by pathologists and compared to patient tumors. Positive staining is expressed as percentage.

### Animals

Non-obese diabetic/severe combined immunodeficiency (NOD/SCID) mice were purchased from Harlan Laboratories. NOD.Cg-Prkdc^scid^ Il2rg^tm1Wjl^/SzJ (NSG) mice were purchased from Charles River. Only female mice 6-12 weeks old (15-20 gr weight) were used for experimental procedures.

### Ethic statements

Investigations have been conducted in accordance with the ethical standards and according to National and International guidelines. *In vivo* studies were performed after approval from our fully authorized animal facility, notification of the experiments to the Ministry of Health (as required by the Italian Law) (IACUCs N° 757/2015, 1246/2015 and 327/2016) and in accordance to EU directive 2010/63. Human tissue biopsies were collected from patients whose informed consent was obtained in writing according to the policies of the Ethics Committee of the European Institute of Oncology and regulations of Italian Ministry of Health. The studies were conducted in full compliance with the Declaration of Helsinki.

### PDX culture

PDX tumors were dissociated by enzymatic and mechanical digestion (Miltenyi Biotec) and cells were plated to obtain short-term cultures. PDX cells were maintained in DMEM/F12 (1:1, Lonza/ Gibco) supplemented with 10% Fetal Bovine Serum (FBS) (HyClone, GE Healthcare Life Science), 10mM HEPES (Sigma), 5 μg/mL insulin (Roche), 0.5 μg/mL hydrocortisone (Sigma), 10 ng/mL epidermal growth factor (EGF, Tebu-Bio), 50 ng/mL Cholera Toxin (Sigma).

### Cell lines

Experiments were performed in: MCF10DCIS.com (from Wayne State University, 5057 Woodward Avenue, Detroit - Michigan), SUM149PT (from Asterand), HCC1428 (from IFOM), MDA-MB-468 (from CLS), ZR751 (from ATCC) and MDA-MB-231 (from NIH Institute) cell lines were maintained in their respective media as recommended by suppliers. Cell line authentication was performed in house by Gene Print 10 System every six months (Promega). All cell lines were tested for mycoplasma and resulted negative.

### *In vivo* study

PDXs, MCF10DCIS.com, SUM149PT, HCC1428, MDAMB468 and ZR751 were infected with control shRNA (shLuc) and pooled shWDR5. 2.5^10^5^ infected cells were orthotopically injected into the 4^th^ mammary gland of 3 to 8 mice (MCF10DCIS.com were transplanted in NOD/SCID mice; PDXs cells, SUM149PT, HCC1428, MDAMB468 AND ZR751 in NSG mice). Tumor volume was calculated using this formula: V=l^2^ΛL/2 (l length; L width). MDA-MB-231 cells were transfected with the luciferase plasmid (Addgene 17477) and selected by using Puromycin. Cells were then transfected to silence shLuc or shWDR5 and then 2^10^5^ transplanted in the 4^th^ mammary gland of 5 NSG mice *per* group. The mice were monitored for primary tumor growth. When a volume of about 0.5 cm^3^ was reached, tumors were excised and mice monitored weekly for metastasis formation. Luciferase expression was assessed by bioluminescence imaging (IVIS Lumina Imaging System - PerkinElmer) and mice were sacrificed one month after resection, when lungs and axillary lymph nodes resulted positive to luminescence. Organs were fixed in paraffin, sectioned and Hematoxylin and eosin (H&E) stained.

### *In vitro* study

Proliferation, migration on Boyden chamber and wound healing assays were performed as described in expanded view.

### Immunofluorescence

MCF10DCIS.com cells were infected to silence shLuc or sWDR5 and after two days of infection 1^10^5^ cells were plated on slides and allowed to attach overnight. Next day, cells were fixed with 4% paraformaldhehyde for 10 minutes, permeabilized with 0.01% Triton-X and blocked for 1h with 2% bovine serum albumine. The actin cytoskeleton was stained using FITC-labeled Phalloidin from Sigma (P5282) for 2 hours. Slides were then counterstained with 4′,6-diamidino-2-phenylindole (DAPI) for nuclei labelling and mounted on glass slides with Mowiol. Images were collected by motorized Olympus fluorescence microscope at 40X magnification.

### Adhesion assay

For adhesion assay, 2^10^4^ shLuc and shWDR5 MCF10DCIS.com cells were plated onto different extracellular matrices (collagen - CL, laminin - LM, fibronectin - FN, matrigel – MG). After 1.5 hour, cells were fixed and stained with 0.5% Crystal Violet. Three images *per* well were acquired and cell number and area were quantified by using ImageJ software by manually delineating the edges of randomly selected cells (a total of 30 measurements *per* group) and recording the circularity value.

### Western blot

PDXs and MCF10DCIS.com cells were lysed in RIPA buffer and processed, as previously described (D’Alesio et al., 2016). Membranes were probed with antibodies reported in expanded. Images were cropped at specific protein band of interest to improve the clarity of data presentation.

### Survival Analysis

Association between WDR5 expression and metastasis free survival (MFS) in 295 breast cancer patients was calculated using PROGgene V2 software on NKI data sets. MFS were represented by Kaplan-Meier functions and cohorts were divided at the median of gene expression. Statistic comparison between high and low expression groups was performed using log rank test. Differences were considered significant at *P* < 0.05.

### RNA-sequencing

Total RNA was extracted from shLuc and shWDR5 MCF10DCIS.com cells or PDXs infected cells by using the Maxwell 16LEV simply RNA tissue kit. mRNA purification and NGS libraries were obtained following Illumina instruction (TruSeq RNA Sample Preparation). Bioinformatic analysis is fully described in Appendix Supplementary Methods.

### ChIP-sequencing

ChIP lysates were generated from 10-15^10^6^ cells as reported previously (Bossi, Cicalese et al., 2016). ChIP DNA was prepared for HiSeq 2000 Illumina sequencing. Samples were aligned to human genome and bioinformatic analysis is fully described in Appendix Supplementary Methods.

## Data Access

Data sets are available in the Gene Expression Omnibus (GEO) database under accession number GSE113289.

### Quantitative RT-PCR

Total RNA was extracted from PDXs and MCF10DCIS.com as above and reverse transcribed using OneScript Plus Reverse Transcriptase and cDNA Synthesis kit (abm). Quantitative RT-PCR analyses were done on Biorad CFX Real-Time PCR System with the fast-SYBR Green PCR kit as instructed by the manufacturer (Applied Biosystems). The transcription level of the RPLP0 housekeeper gene was used as normalizer. Complete primers sequences are reported in Appendix Table S3.

### Drugs treatment

1.5^10^6^ cells were plated on 10cm plates. Recombinant human TGFβ1 (Peprotech) was added at a final concentration of 5ng/ml for 2 days. OICR-9429 was obtained from MD Anderson Cancer Center (Texas) and resuspended in DMSO. 1.5^10^5^ MCF10DCIS.com were plated in six-well plates. Drug was added to complete medium at a final concentration of 1μM - 5μM - 10μM or 20μM for 3 days and cells were then effects on proliferation was assessed. 10^3^ cells were plated for colony formation assay and drug was added for 6 days (medium and drug was refreshed on day 3). Cells were then stained with Crystal violet and quantified by ImageJ analysis.

### Statistical analysis

Data are represented as mean ± SD of triplicate biological replicates (if not diversely indicated in the text) and have been statistically assessed by two-tailed Student’s *t* test as indicate in figure legend. In figures, asterisks mean * p<0.05, ** p<0.01, *** p<0.001. p<0.05 and lower were considered significant. For RNA-seq and ChIP-seq analysis, statistical parameters are indicated in Appendix Supplementary methods.

## Acknowledgments

S. Punzi is recipient of a FUV fellowship. This work was supported by the European Research Council Advanced Grant 341131. We thank A. De Rose and R. Piccioni for technical support. We also thank A. Gobbi and M. Capillo for excellent support in animal work. We wish to thank all members of the Department of Experimental Oncology for discussion and reagents. We thank the Genomic Unit (IEO), the Mouse Facility (Cogentech), Istituto Italiano di Tecnologia (IIT) and the Cell Biology Unit (IEO).

## Author contribution

S.P., L.L., conceived the study, designed the experiments and interpreted the data. C.B. performed computational analysis. C.D., D.B., A.C., G.I.D. and E.G. provided essential reagents and expertise. G.P., C.C. and G.C. managed patients. P.G.P, G.N. reviewed critically the work. L.L. supervised the work. S.P. and L.L. wrote the manuscript.

## Expanded view Figure Legend

**Figure EV 1.**
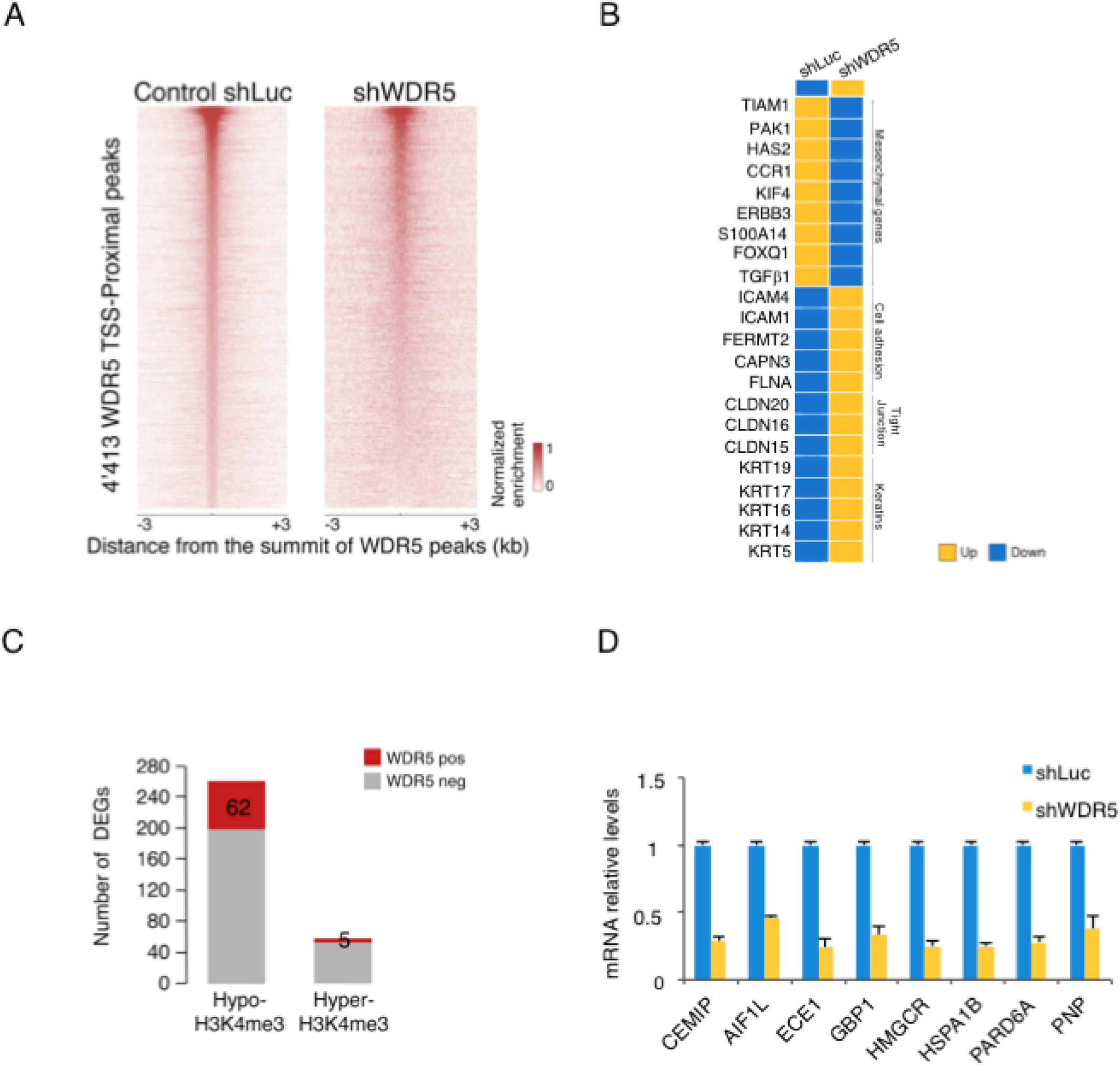
WDR5 regulates TGFβ targets. A) Heatmap of Transcription Start Sites (TSS)-proximal WDR5 binding events in the shLuc MCF10DCIS cancer cell line is shown. A 3kb genomic interval centered on the summit of WDR5 peaks is considered. For comparison, the same regions are also shown in shWDR5 MCF10DCIS cells. B) Heatmap represents down-(orange) and up-(blue) regulated genes involved in epithelial and mesenchymal control in MCF10DCIS cell line due to WDR5 silencing. C) DEGs within differentially methylated promoters are shown. Genes with differential expression and differential H3K4me3 but not bound by WDR5 are in gray and genes with differential expression, differential H3K4me3 and bound WDR5 peaks are in red. D) Relative expression of main WDR5 targets highlighted by RNA-seq analysis was detected in shWDR5 MCF10DCIS cells by RT-qPCR with respect to shLuc. The results are presented as means ± SD of three independent experiments.

### Expanded view Table Legend

**Table EV 1.** WDR5 genome wide binding sites on MCF10DCIS were assigned to the nearest proximal and distal transcription start sites (TSS)(+3kb). H3K4me3 ChIP-seq was performed on MCF10DCIS breast cancer cell line. Reads were mapped to the promoter region (±1’500 bp relative to TSS) for annotated transcripts of DEGs. Significant differential H3K4me3 values are shown.

